# Acute myeloid/T-lymphoblastic leukemia (AMTL): A distinct category of acute leukemias with common pathogenesis in need of improved therapy

**DOI:** 10.1101/223388

**Authors:** Alejandro Gutierrez, Alex Kentsis

**Author notes:** Correspondence to: Alejandro Gutierrez, Alex Kentsis.

## Abstract

Advances in the immunophenotypic and cytogenetic classification of acute leukemias have led to improved clinical outcomes for a substantial fraction of patients. However, resistance to chemotherapy remains a major barrier to cure for patients with specific subsets of acute myeloid and lymphoblastic leukemias. Here, we propose that a molecularly distinct subtype of acute leukemia with shared myeloid and T-cell lymphoblastic features, which we term acute myeloid/T-lymphoblastic leukemia (AMTL), and has been divided between 3 diagnostic categories owing to variable expression of markers deemed to be defining of myeloid and T-cell lymphoid lineages. This new diagnostic group is supported by the i) shared hematopoietic ontogeny in which myeloid differentiation potential is specifically retained during early T-cell lymphoid development, ii) recognition of cases of AML with hallmarks of T-cell development such as clonal rearrangements of the T-cell receptor genes, and iii) identification of common gene mutations in subsets of AML and T-ALL cases. This proposed diagnostic entity overlaps with early T-cell precursor (ETP) T-ALL and T-cell/myeloid mixed phenotype acute leukemias (MPAL), and also includes a subset of leukemias currently classified as AML with hallmarks of T-lymphoblastic development. AMTLs express variable levels of both T-cell and myeloid-specific markers, such as CD3 and myeloperoxidase, and additionally have shared gene mutations including *WT1, PHF6, RUNX1* and *BCL11B*. The proposed classification of AMTL as a distinct entity should enable prospective diagnosis and development of improved therapies for patients whose treatment is inadequate with current approaches.

## Introduction

Acute leukemias are aggressive neoplasms characterized by the pathologic accumulation of immature hematopoietic progenitors. Improvements in clinical outcomes for patients with acute leukemias treated with modern chemotherapy regimens have been uneven. The introduction of molecularly targeted therapies has had a major impact on a small subset of genetically defined acute leukemias. For example, arsenic trioxide and retinoic acid, which target the PML-RARA fusion oncoprotein via distinct mechanisms (1), can now cure the majority of patients with acute promyelocytic leukemia without exposure to cytotoxic chemotherapies (2). However, clinical outcomes for a substantial fraction of acute leukemias have seen little improvement despite the application of treatment regimens that are among the most intensive and toxic used for any disease. Consequently, improving clinical outcomes for these patients will require linking an improved molecular understanding to the application of effective targeted therapies.

### Classification of acute leukemias

Acute leukemias have traditionally been classified based on the normal cell types most closely resembling the leukemic cell population. Lymphoblastic leukemias are those with evidence of differentiation arrest at immature stages of B- or T-cell lymphoid development, whereas acute myeloid leukemias encompass malignancies with immunophenotypic features that lie along a broad spectrum of hematopoietic progenitors, ranging from minimally differentiated leukemias to those with evidence of granulocytic, monocytic, erythroid or megakaryocytic differentiation. Although this classification scheme is intuitive, lineage assignment is complicated by the high frequency of aberrant differentiation states in acute leukemias. Indeed, the simultaneous expression of markers that are not known to be coexpressed in any normal hematopoietic progenitor is common. To address the requirement for consistent diagnostic criteria for the successful performance and interpretation of clinical trials, the World Health Organization (WHO) has devised a classification scheme that allows most acute leukemias to be unambiguously classified as specific diagnostic entities, as regularly revised (3, 4).

The genetics of leukemia are beginning to be incorporated into the WHO classification, with increasing weight being given to specific chromosomal translocations, and in some cases, somatic gene mutations, as disease-defining genetic lesions (3). Thus, the presence of a characteristic genetic feature can define a disease, independent of the phenotypic lineage which can be difficult to assign in practice. For example, leukemias with FGFR1 translocations that can present with T-cell lymphoblastic or myeloid markers are defined as a single diagnostic entity, regardless of the immunophenotype of the presenting leukemic cells (3, 5). Similarly, AML cases with t(8;21) translocations exhibit aberrant expression of CD19, a marker typically associated with B-cell lymphoid malignancies, due to lineage-inappropriate PAX5 expression induced by the AML1-ETO oncogene (6). While we anticipate that genomic classification will have an increasingly important role in defining leukemia subsets in the future, the current diagnostic classification for most acute leukemias does not rely on genetic mutations. Thus, phenotypic lineage assignment remains a central component of diagnostic classification and treatment assignment for most patients with acute leukemia.

The current WHO classification considers high expression of a small number of immunophenotypic markers to be lineage-assigning: myeloperoxidase (MPO) or at least two monocytic markers (CD11c, CD14, CD64, or lysozyme) for myeloid lineage; surface or cytoplasmic CD3 expression for T-cell lineage; and CD19 with at least one additional B-cell marker (CD79a, cytoplasmic CD22, or CD10) for B-cell lineage. However, it is recognized that these criteria are imperfect, and their strict application is not required to make a diagnosis of AML, T-ALL or B-ALL unless a diagnosis of mixed phenotype acute leukemia (MPAL) is also being considered (3). Thus, while mixed phenotype acute leukemias that meet these strict criteria for assignment to two distinct lineages are very rare, leukemias classified as AML, T-ALL or B-ALL based on their predominant morphology and immunophenotype commonly co-express markers that indicate aberrant differentiation towards another lineage.

### Acute myeloid/T-lymphoblastic leukemia (AMTL): Acute leukemias with shared T-cell lymphoid and myeloid features

The classical model of hematopoiesis postulates an early binary split between common myeloid and lymphoid progenitors that subsequently give rise to both B- and T-cell lymphocytes. However, more recent work using clonal tracking assays has shown the existence of progenitors that retain T-cell/myeloid or B-cell/myeloid bi-lineage potential, whereas individual progenitors whose potential is restricted to T- and B-cell lymphoid lineages have been difficult to identify (7). These findings fit the clinical observation that mixed phenotype acute leukemias most commonly present with B-cell/myeloid or T-cell/myeloid marker co-expression, whereas B/T-cell mixed phenotype acute leukemias are very rare (8).

Here, we propose AMTL as a molecularly distinct subtype of acute leukemias associated with shared T-cell lymphoid and myeloid features. Expression of markers deemed to be defining of the myeloid and T-cell lymphoid lineages, including myeloperoxidase and CD3, is variable in this disease, resulting in its separation across three diagnostic categories in the current WHO classification scheme: ETP T-ALL, T/myeloid mixed phenotype acute leukemia (MPAL), and a specific subset of AML harboring hallmarks of T-lymphoblastic differentiation.

This proposed diagnostic entity overlaps with subsets of early T-cell precursor (ETP) TALL, which is characterized by differentiation arrest at early stages of T-cell development. While a number of biomarkers have been described to identify these cases (9–12), these are most commonly defined clinically using an immunophenotypic classifier that includes absent expression of CD1a and CD8, expression of CD5 that is substantially lower than that of normal peripheral blood T-cells, and the presence of one or more markers of myeloid or hematopoietic progenitors (CD117, CD34, HLA-DR, CD13, CD33, CD11b or CD65) (9). ETP is classified as a subtype of T-ALL due to the expression of CD3, but has a genetic mutational profile that is similar to those observed in myeloid malignancies such as AML, whereas mutations characteristic of other subtypes of T-ALL, such as *CDKN2A* deletions, are less common (11, 13, 14). Whether T-ALL cases expressing the ETP immunophenotype that harbor classical T-ALL oncogenic lesions, such as *NOTCH1* mutations, are molecularly distinct from ETPs with a more typically myeloid-like mutational signature remains to be defined.

While similarities between ETP T-ALL and AML have been previously recognized (10, 13, 14), we propose that the biologic overlap is not with AML per se, but with a specific subset of AML cases that also exhibit T-cell lymphoid features. Namely, a subset of AMLs has long been recognized to harbor clonal T-cell receptor (TCR) or immunoglobulin (Ig) gene rearrangements, indicating activity of the RAG recombinase that is responsible for generating somatic V(D)J recombination at these loci at specific stages of lymphoid development (15–17). These AML cases can also express the lymphoid marker terminal deoxynucleotidyltransferase (TdT, also known as DNTT), which generates diversity at TCR and Ig genes by mutating the junctions of rearrangements during V(D)J recombination (18, 19). Non-megakaryocytic AMLs with T-cell receptor rearrangements typically co-express phenotypic markers of T-lymphoblastic differentiation, such as CD7, CD2 and CD4 (16). We postulate that this is the subset of AMLs that also harbor mutations that are commonly observed in a distinct subset of T-ALL, including *WT1, PHF6, RUNX1* and *BCL11B* (13, 20–26). Thus, a similar group of acute leukemias exhibit shared features of myeloid and T-cell lymphoid differentiation, and shared genetic mutations, which will need to be formally assessed in future studies.

### Potential mechanisms for combined T-lymphoblastic and myeloid differentiation in acute leukemia

At least two non-mutually exclusive mechanisms can explain the development of acute leukemias with differentiation potential restricted to myeloid and T-lymphoid lineages:

1. *Transformation of normal hematopoietic progenitors with T-lymphoblastic and myeloid bi-lineage potential*. While studies of endogenous hematopoiesis have revealed limited evidence of a putative normal progenitor whose fate is restricted to T-lymphoid and myeloid lineages, the identification of such a progenitor may have been hindered by the differences in lifespan of a mature myeloid versus T-lymphoid cells. Nevertheless, even if a normal progenitor whose differentiation fate is restricted to T-lymphocytes and myeloid cells is lacking, such leukemias can arise from cells that retain multi-lineage potential, even though their fate is normally restricted to a single lineage by microenvironmental signals. Immature intrathymic T-cell progenitors at early double-negative stages of differentiation represent one potential cell of origin for such leukemias. Indeed, the most immature intrathymic T-cell progenitors retain myeloid (but not B-lymphoid) potential until they progress through a BCL11B-dependent commitment to the T-cell lineage (27–29) (Figure 1). Further, in vivo lineage tracking studies have demonstrated that immature T-cell progenitors give rise to some macrophages and neutrophils harboring T-cell receptor gene rearrangements in vivo (30, 31). Ectopic expression of MYC and BCL2 in mouse early double-negative T-cell progenitors can drive development of acute leukemias with variable expression of markers of myeloid and T-lymphoid lineages (32), demonstrating that early T-cell progenitors can function as the cell of origin of AMTL.
2. *Induction of aberrant trans-differentiation by leukemogenic mutations*. It is now established that specific gene mutations can induce T-cell lymphoid or myeloid differentiation. For example, activation of Notch1 signaling in mouse bone marrow cells causes aberrant T-cell differentiation independent of thymic microenvironmental signals (33). Likewise, mutations inactivating *Runx1* in hematopoietic progenitors causes aberrant myeloid differentiation (34). Thus, aberrant trans-differentiation of AMTL cells can in principle occur as a result of combinations of mutations that cooperatively result in combined myeloid and T-lymphoid lineage commitment with differentiation arrest (Figure 1). These mutations include genes encoding regulators of chromatin and DNA remodeling, and transcription factors with key functions in controlling lymphoid and myeloid cell fate specification. Future studies using emerging approaches for combinatorial gene editing and conditional gene manipulation in vivo may reveal how the specific combinations of mutations in specific cell progenitor populations cooperate to induce AMTL and other non-canonical leukemia subtypes.

**Figure 1.**
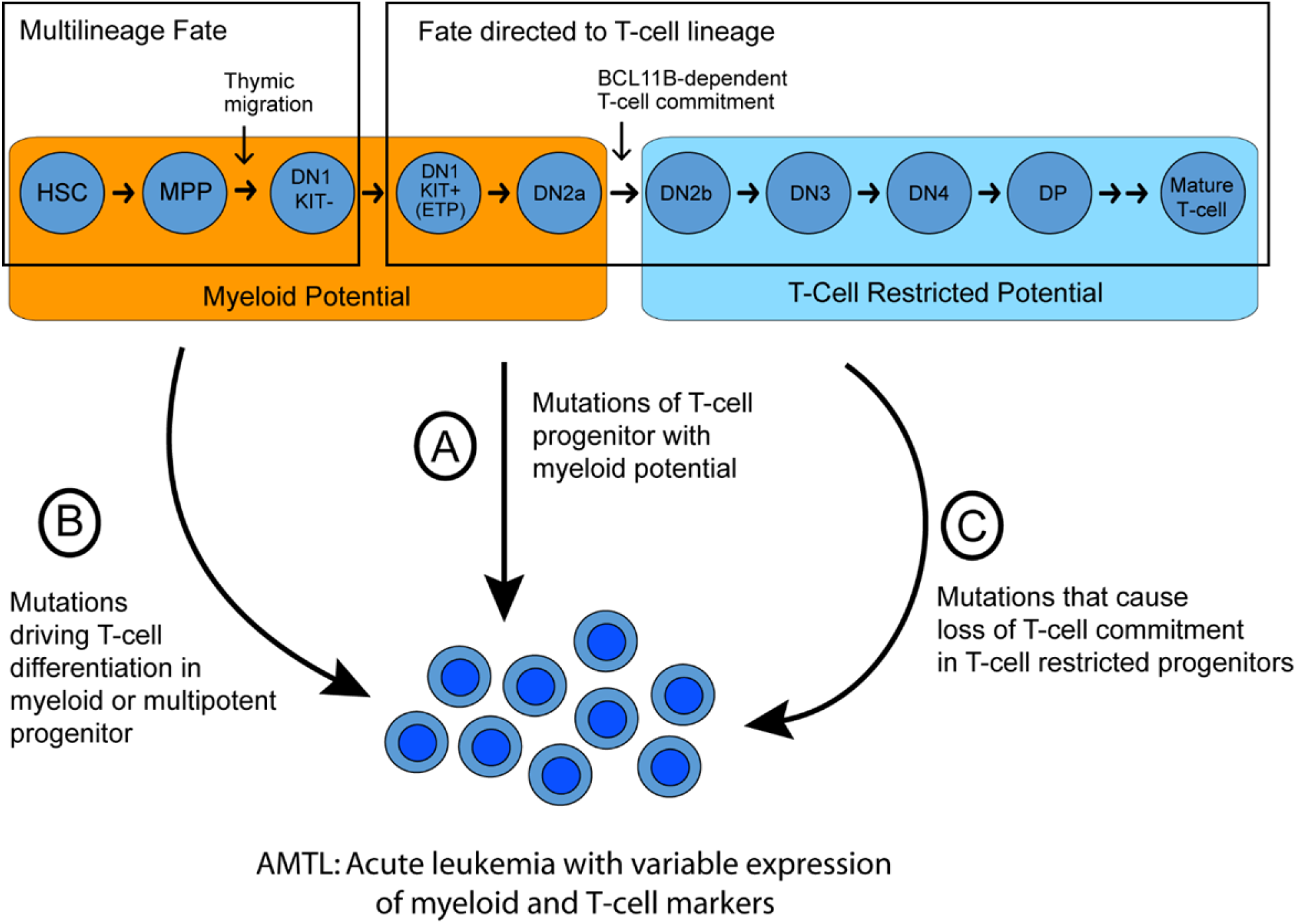
Potential mechanisms leading to acute leukemia with shared myeloid and T-lymphoblastic features. The fate of multipotent hematopoietic progenitors first entering the thymus is directed to the T-cell lineage via thymic microenvironmental signals, but these cells retain myeloid potential until a BCL11B-dependent commitment to the T-cell lineage at the DN2a to DN2b transition (40). Such T-cell progenitors with myeloid potential can function as a cell of origin of AMTL (32) (A). Alternatively, AMTL can in principle arise from mutations that induce aberrant T-cell differentiation in myeloid progenitors (B), or from mutations in T-cell restricted progenitors leading to their myeloid differentiation (C). Note that the developmental stages shown (Top) are based on mouse T-cell development, because human early T-cell development is less well defined. *HSC, hematopoietic stem cell. MPP, multipotent progenitor. DN, CD4/CD8 double-negative T-cell progenitor. DP, CD4/CD8 double-positive progenitor*.

### Identification of Acute Myeloid/T-Lymphoblastic Leukemia

To a first approximation, AMTL can be defined as acute leukemias that fit the diagnostic criteria of ETP T-ALL or of T/myeloid mixed phenotype acute leukemias, together with AMLs with clonal T-cell receptor gene rearrangements and evidence of T-lymphoid differentiation (CD3, CD7, CD2 or CD4 expression). However, we note that aberrant expression of CD4 and CD7 is common in acute megakaryoblastic leukemia, a subtype of AML that is molecularly distinct from AMTL (35). Thus, acute megakaryoblastic leukemia should be excluded from the definition of AMTL, based on absence of megakaryblastic morphology and expression of the platelet markers CD41 or CD61.

Given the variability in cell surface marker expression in AMTL, we propose that clinical genomic profiling will enhance the classification of AMTL as a specific diagnostic entity. Indeed, ETP T-ALLs harbor mutations of several genes that are commonly mutated in myeloid neoplasms, such as RAS, ETV6, MEF2C and EZH2, but whose mutations are rare in other TALL subtypes, (10, 12–14). We also suggest that the subset of AML cases harboring clonal T-cell receptor gene rearrangements should demonstrate substantial overlap with those AML cases harboring mutations that are common in T-ALL, such as WT1, *RUNX1, PHF6*, and *BCL11B* (13, 20–22). Thus, we anticipate that further investigation will lead to an improved classifier of AMTL based on both immunophenotype and genetics.

## Conclusions

We expect that defining the detailed molecular mechanisms of aberrant cell differentiation may lead to the development of improved therapies by targeting specific molecular dependencies in these cells. For example, mutant transcription factors function in the context of corepressor and coactivator complexes, which have specific enzymatic functions, such as histone deacetylase (HDAC) inhibitors (36). Isoform-specific HDAC inhibitors are beginning to be developed (37), and specific inhibitors may warrant therapeutic investigation in AMTL with dysregulation of MEF2 family members, for example. Likewise, specific mutations may engender synthetic lethal dependencies, such as for example inhibition of the EZH2 methyltransferase in cases of cancers with mutations of genes encoding components of the SWI/SNF/BAF chromatin remodeling complex (38). Recently, acetyltranferase inhibitors have been found to have therapeutic efficacy in CBP-deficient lymphomas (39). Finally, many leukemias exhibit aberrant resistance to mitochondrial apoptosis, conferring resistance to intensive combination chemotherapies, such as those used for treatment of refractory AML and T-ALL. Emerging inhibitors of BCL2 and other regulators of intrinsic apoptosis are expected to offer useful therapeutic strategies for AMTL patients as well.

## Acknowledgements

We thank Misha Roshal, Mark Fleming, and Peter Steinherz for helpful discussions. This work was supported by the NIH R01 CA204396 and CA193651, and P30 CA008748. A.G. and A.K. are Damon Runyon Clinical Investigators.

## References

1. de The H. Lessons taught by acute promyelocytic leukemia cure. Lancet. 2015;386(9990):247–8.

2. Lo-Coco F, Avvisati G, Vignetti M, Thiede C, Orlando SM, Iacobelli S, et al. Retinoic acid and arsenic trioxide for acute promyelocytic leukemia. N Engl J Med. 2013;369(2):111–21.

3. Arber DA, Orazi A, Hasserjian R, Thiele J, Borowitz MJ, Le Beau MM, et al. The 2016 revision to the World Health Organization classification of myeloid neoplasms and acute leukemia. Blood. 2016;127(20):2391–405.

4. WHO Classification of Tumours of Haematopoietic and Lymphoid Tissues. Swerdlow SH, Campo E, Harris NL, Jaffe ES, Pileri SA, Stein H, et al., editors. Lyon: IARC Press; 2008.

5. Macdonald D, Reiter A, Cross NC. The 8p11 myeloproliferative syndrome: a distinct clinical entity caused by constitutive activation of FGFR1. Acta Haematol. 2002;107(2):101–7.

6. Walter K, Cockerill PN, Barlow R, Clarke D, Hoogenkamp M, Follows GA, et al. Aberrant expression of CD19 in AML with t(8;21) involves a poised chromatin structure and PAX5. Oncogene. 2010;29(20):2927–37.

7. Kawamoto H, Ikawa T, Masuda K, Wada H, Katsura Y. A map for lineage restriction of progenitors during hematopoiesis: the essence of the myeloid-based model. Immunol Rev. 2010;238(1):23–36.

8. Matutes E, Pickl WF, Van’t Veer M, Morilla R, Swansbury J, Strobl H, et al. Mixed-phenotype acute leukemia: clinical and laboratory features and outcome in 100 patients defined according to the WHO 2008 classification. Blood. 2011;117(11):3163–71.

9. Coustan-Smith E, Mullighan CG, Onciu M, Behm FG, Raimondi SC, Pei D, et al. Early T-cell precursor leukaemia: a subtype of very high-risk acute lymphoblastic leukaemia. Lancet Oncol. 2009;10(2):147–56.

10. Zuurbier L, Gutierrez A, Mullighan CG, Cante-Barrett K, Gevaert AO, de Rooi J, et al. Immature MEF2C-dysregulated T-cell leukemia patients have an early T-cell precursor acute lymphoblastic leukemia gene signature and typically have non-rearranged T-cell receptors. Haematologica. 2014;99(1):94–102.

11. Gutierrez A, Dahlberg SE, Neuberg DS, Zhang J, Grebliunaite R, Sanda T, et al. Absence of biallelic TCRgamma deletion predicts early treatment failure in pediatric T-cell acute lymphoblastic leukemia. J Clin Oncol. 2010;28(24):3816–23.

12. Homminga I, Pieters R, Langerak AW, de Rooi JJ, Stubbs A, Verstegen M, et al. Integrated transcript and genome analyses reveal NKX2-1 and MEF2C as potential oncogenes in T cell acute lymphoblastic leukemia. Cancer Cell. 2011;19(4):484–97.

13. Zhang J, Ding L, Holmfeldt L, Wu G, Heatley SL, Payne-Turner D, et al. The genetic basis of early T-cell precursor acute lymphoblastic leukaemia. Nature. 2012;481(7380):157–63.

14. Van Vlierberghe P, Ambesi-Impiombato A, Perez-Garcia A, Haydu JE, Rigo I, Hadler M, et al. ETV6 mutations in early immature human T cell leukemias. J Exp Med. 2011;208(13):2571–9.

15. Norton JD, Campana D, Hoffbrand AV, Janossy G, Coustan-Smith E, Jani H, et al. Rearrangement of immunoglobulin and T cell antigen receptor genes in acute myeloid leukemia with lymphoid-associated markers. Leukemia. 1987;1(11):757–61.

16. Schmidt CA, Oettle H, Neubauer A, Seeger K, Thiel E, Huhn D, et al. Rearrangements of T-cell receptor delta, gamma and beta genes in acute myeloid leukemia coexpressing T-lymphoid features. Leukemia. 1992;6(12):1263–7.

17. Parreira L, Carvalho C, Moura H, Melo A, Santos P, Guimaraes JE, et al. Configuration of immunoglobulin and T cell receptor beta and gamma genes in acute myeloid leukaemia: pitfalls in the analysis of 40 cases. J Clin Pathol. 1992;45(3):193–200.

18. Drexler HG, Sperling C, Ludwig WD. Terminal deoxynucleotidyl transferase (TdT) expression in acute myeloid leukemia. Leukemia. 1993;7(8):1142–50.

19. Patel KP, Khokhar FA, Muzzafar T, James You M, Bueso-Ramos CE, Ravandi F, et al. TdT expression in acute myeloid leukemia with minimal differentiation is associated with distinctive clinicopathological features and better overall survival following stem cell transplantation. Mod Pathol. 2013;26(2):195–203.

20. Cancer Genome Atlas Research N, Ley TJ, Miller C, Ding L, Raphael BJ, Mungall AJ, et al. Genomic and epigenomic landscapes of adult de novo acute myeloid leukemia. N Engl J Med. 2013;368(22):2059–74.

21. Klco JM, Miller CA, Griffith M, Petti A, Spencer DH, Ketkar-Kulkarni S, et al. Association Between Mutation Clearance After Induction Therapy and Outcomes in Acute Myeloid Leukemia. JAMA. 2015;314(8):811–22.

22. Ding L, Ley TJ, Larson DE, Miller CA, Koboldt DC, Welch JS, et al. Clonal evolution in relapsed acute myeloid leukaemia revealed by whole-genome sequencing. Nature. 2012;481(7382):506–10.

23. Tosello V, Mansour MR, Barnes K, Paganin M, Sulis ML, Jenkinson S, et al. WT1 mutations in T-ALL. Blood. 2009;114(5):1038–45.

24. Van Vlierberghe P, Patel J, Abdel-Wahab O, Lobry C, Hedvat CV, Balbin M, et al. PHF6 mutations in adult acute myeloid leukemia. Leukemia. 2011;25(1):130–4.

25. Van Vlierberghe P, Palomero T, Khiabanian H, Van der Meulen J, Castillo M, Van Roy N, et al. PHF6 mutations in T-cell acute lymphoblastic leukemia. Nat Genet. 2010;42(4):338–42.

26. Della Gatta G, Palomero T, Perez-Garcia A, Ambesi-Impiombato A, Bansal M, Carpenter ZW, et al. Reverse engineering of TLX oncogenic transcriptional networks identifies RUNX1 as tumor suppressor in T-ALL. Nat Med. 2012;18(3):436–40.

27. Li L, Leid M, Rothenberg EV. An early T cell lineage commitment checkpoint dependent on the transcription factor Bcl11b. Science. 2010;329(5987):89–93.

28. Ikawa T, Hirose S, Masuda K, Kakugawa K, Satoh R, Shibano-Satoh A, et al. An essential developmental checkpoint for production of the T cell lineage. Science. 2010;329(5987):93–6.

29. Balciunaite G, Ceredig R, Rolink AG. The earliest subpopulation of mouse thymocytes contains potent T, significant macrophage, and natural killer cell but no B-lymphocyte potential. Blood. 2005;105(5):1930–6.

30. Bell JJ, Bhandoola A. The earliest thymic progenitors for T cells possess myeloid lineage potential. Nature. 2008;452(7188):764–7.

31. Wada H, Masuda K, Satoh R, Kakugawa K, Ikawa T, Katsura Y, et al. Adult T-cell progenitors retain myeloid potential. Nature. 2008;452(7188):768–72.

32. Riemke P, Czeh M, Fischer J, Walter C, Ghani S, Zepper M, et al. Myeloid leukemia with transdifferentiation plasticity developing from T-cell progenitors. EMBO J. 2016;35(22):2399–416.

33. Pui JC, Allman D, Xu L, DeRocco S, Karnell FG, Bakkour S, et al. Notch1 expression in early lymphopoiesis influences B versus T lineage determination. Immunity. 1999;11(3):299–308.

34. Goyama S, Schibler J, Cunningham L, Zhang Y, Rao Y, Nishimoto N, et al. Transcription factor RUNX1 promotes survival of acute myeloid leukemia cells. J Clin Invest. 2013;123(9):3876–88.

35. de Rooij JD, Branstetter C, Ma J, Li Y, Walsh MP, Cheng J, et al. Pediatric non-Down syndrome acute megakaryoblastic leukemia is characterized by distinct genomic subsets with varying outcomes. Nat Genet. 2017;49(3):451–6.

36. Haberland M, Montgomery RL, Olson EN. The many roles of histone deacetylases in development and physiology: implications for disease and therapy. Nat Rev Genet. 2009;10(1):32–42.

37. West AC, Johnstone RW. New and emerging HDAC inhibitors for cancer treatment. J Clin Invest. 2014;124(1):30–9.

38. Kim KH, Kim W, Howard TP, Vazquez F, Tsherniak A, Wu JN, et al. SWI/SNF-mutant cancers depend on catalytic and non-catalytic activity of EZH2. Nat Med. 2015;21(12):1491–6.

39. Ogiwara H, Sasaki M, Mitachi T, Oike T, Higuchi S, Tominaga Y, et al. Targeting p300 Addiction in CBP-Deficient Cancers Causes Synthetic Lethality by Apoptotic Cell Death due to Abrogation of MYC Expression. Cancer Discov. 2016;6(4):430–45.

40. Hosokawa H, Rothenberg EV. Cytokines, Transcription Factors, and the Initiation of T-Cell Development. Cold Spring Harb Perspect Biol. 2017.

